# A consensus set of genetic vulnerabilities to ATR inhibition

**DOI:** 10.1101/574533

**Authors:** Nicole Hustedt, Alejandro Álvarez-Quilón, Andrea McEwan, Jing Yi Yuan, Tiffany Cho, Lisa Koob, Traver Hart, Daniel Durocher

## Abstract

The response to DNA replication stress in eukaryotes is under the control of the ataxia-telangiectasia and Rad3-related (ATR) kinase. ATR responds to single-stranded (ss) DNA to stabilize distressed DNA replication forks, modulate DNA replication firing and prevent cells with damaged DNA or incomplete DNA replication from entering into mitosis. Furthermore, inhibitors of ATR are currently in clinical development either as monotherapies or in combination with agents that perturb DNA replication. To gain a genetic view of the cellular pathways requiring ATR kinase function, we mapped genes whose mutation causes hypersensitivity to ATR inhibitors with genome-scale CRISPR/Cas9 screens. We delineate a consensus set of 117 genes enriched in DNA replication, DNA repair and cell cycle regulators that promote survival when ATR kinase activity is suppressed. We validate 14 genes from this set and report genes not previously described to modulate response to ATR inhibitors. In particular we found that the loss of the POLE3/POLE4 proteins, which are DNA polymerase e accessory subunits, results in marked hypersensitivity to ATR inhibition. We anticipate that this 117-gene set will be useful for the identification of genes involved in the regulation of genome integrity, the characterization of new biological processes involving ATR, and may reveal biomarkers of ATR inhibitor response in the clinic.

## Introduction

The ATR kinase is a phosphoinositide 3-kinase-like kinase that is activated when single-stranded (ss) DNA bound by the RPA complex is sensed by pathways anchored by the ATRIP or ETAA1 proteins (Zou and Elledge 2003; Bass et al. 2016; Feng et al. 2016; Haahr et al. 2016; Lee et al. 2016; Zou 2017). Impaired replication or stalled replisomes often produce DNA structures that contain ssDNA that are then sensed by ATR (Sogo et al. 2002; Zou and Elledge 2003; Byun et al. 2005) Accordingly, ATR is a key modulator of DNA replication where it plays multiple roles in ensuring the orderly execution of DNA synthesis and its coordination with G2 phase entry (Saldivar et al. 2017; Saldivar et al. 2018). Perhaps the best-characterized function of ATR is its role in controlling the timely activation of cyclin-dependent kinases (CDKs) via its activation of the CHK1 kinase (Guo et al. 2000; Liu et al. 2000). The activated ATR-CHK1 pathway suppresses CDK activity by inactivating the phosphatases of the CDC25 family, which are CDK activators (Mailand et al. 2000). A second key role for ATR concerns its function in promoting the stability of distressed replication forks (Lopes et al. 2001; Cobb et al. 2003). ATR impacts replication fork stability at multiple levels, for example by modulating fork reversal via the phosphorylation of proteins such as the annealing helicase SMARCAL1 (Couch et al. 2013), controlling the supply of dNTPs (Zhang et al. 2009; Buisson et al. 2015) and regulating the availability of RPA, which protects ssDNA from unscheduled nucleolysis (Toledo et al. 2013). ATR also controls DNA replication origin firing on both local and global scales (Yekezare et al. 2013; Saldivar et al. 2017). While ATR activation is known to suppress late origin firing in both unperturbed and challenged S phase (Santocanale and Diffley 1998; Costanzo et al. 2003; Karnani and Dutta 2011), origins located in the vicinity of a blocked replication fork are shielded from this global inhibition, resulting in an increase in local origin firing that promotes the completion of DNA synthesis and the rescue of stalled replication forks (Blow and Ge 2009; Ge and Blow 2010; Petermann et al. 2010; Karnani and Dutta 2011).

These critical functions of ATR in coordinating the response to replication stress has made it an attractive therapeutic target in oncology given the observation that tumors often display signs of replication stress (Gaillard et al. 2015; Lecona and Fernandez-Capetillo 2018). Multiple clinical-stage ATR inhibitors (ATRi) are now being tested in cancer treatment either as monotherapies or in combination with other agents (O’Connor 2015; Lecona and Fernandez-Capetillo 2018). While ATRi are showing single-agent efficacy in some patients, there is currently a paucity of robust biomarkers of responses, hampering development of this class of agents. Nevertheless, mutations in *ATM* or *ARID1A*, as well as overexpression of *APOBEC3A/B* overexpression or rearrangements of the *EWSR1, MLL* and *SS18-SSX* genes have all been proposed as candidate patient-selection markers for ATRi clinical development (Reaper et al. 2011; Morgado-Palacin et al. 2016; Nieto-Soler et al. 2016; Williamson et al. 2016; Buisson et al. 2017; Green et al. 2017; Jones et al. 2017).

We reasoned that the unbiased identification of genes promoting viability following ATR inhibition would be useful for two purposes. First, this list may contain genes that have not been previously associated with the regulation of DNA replication, cell cycle progression or DNA repair and may reveal new facets of the function of ATR in promoting genome integrity. Secondly, this gene list may assist in the development of new patient-selection hypotheses or may reveal new genetic markers of ATRi response. Prior to the advent of CRISPR-based genetic screens, the search for genetic interactions with ATR deficiency involved studies in genetically-tractable organisms like budding yeast or the use of RNA interference. For example, a focused screen for synthetic lethal interactions with a partially defective budding yeast ATR mutant, *mec1-100*, identified mutations in genes coding for chromatin remodelers, nuclear envelope components and various transcription regulators (Hustedt et al. 2014). In another example, a focused siRNA screen in human cells surveying 240 DNA repair and replication genes identified deficiency of XPF-ERCC1 as well as knockdown of replication-related genes as conditions that induce ATRi sensitivity (Mohni et al. 2014; Mohni et al. 2015). However, the advent of CRISPR/Cas9-based chemogenomic screens now allow the identification of vulnerabilities to ATR inhibition at the genome-scale level and in a robust manner, which was not previously possible with techniques such as RNA interference.

Therefore, we undertook four genome-scale CRISPR/Cas9 screens and combined their results with those of three additional, recently published screens (Wang et al. 2018). From these 7 screens, performed in five different cell lines and using two different ATRi, we describe a core set of 117 genes that promote cellular resistance to ATR inhibition. In particular, we found that loss-of-function mutations in the genes coding for the POLE3/POLE4 histone-fold complex causes ATRi hypersensitivity in human cells. POLE3 and POLE4 form an ancillary subunit complex of DNA polymerase ε involved in histone deposition at the replication fork (Bellelli et al. 2018a; Yu et al. 2018). Our results using a *POLE3* separation-of-function mutant suggest, however, that impaired histone deposition does not underlie the observed ATRi sensitivity pointing rather to its function in DNA synthesis. We believe that this consensus genetic map of vulnerabilities to ATR inhibition will provide a useful resource to those interested in ATR function and therapeutics.

## Results and Discussion

To identify genes and cellular processes that require ATR kinase activity for cellular fitness, we undertook a set of four CRISPR/Cas9 somatic genetic screens in human cells. The screens, schematically depicted in Fig. 1A, were carried out essentially as described before (Hart et al. 2015; Zimmermann et al. 2018). They entailed the transduction of Cas9-expressing cells with a lentiviral library of single-guide (sg)RNAs and after selection and time for editing, the resulting pool of gene-edited cells was split in two populations. One control population was left untreated for the duration of the screen while a second population was incubated with a sublethal dose of an ATR inhibitor that killed ∼20% of cells. sgRNA abundance was determined in each population after 12 d of treatment by sequencing and a gene-based depletion score was determined with the most up-to-date version of drugZ (Wang et al. 2017).

**Figure 1.**
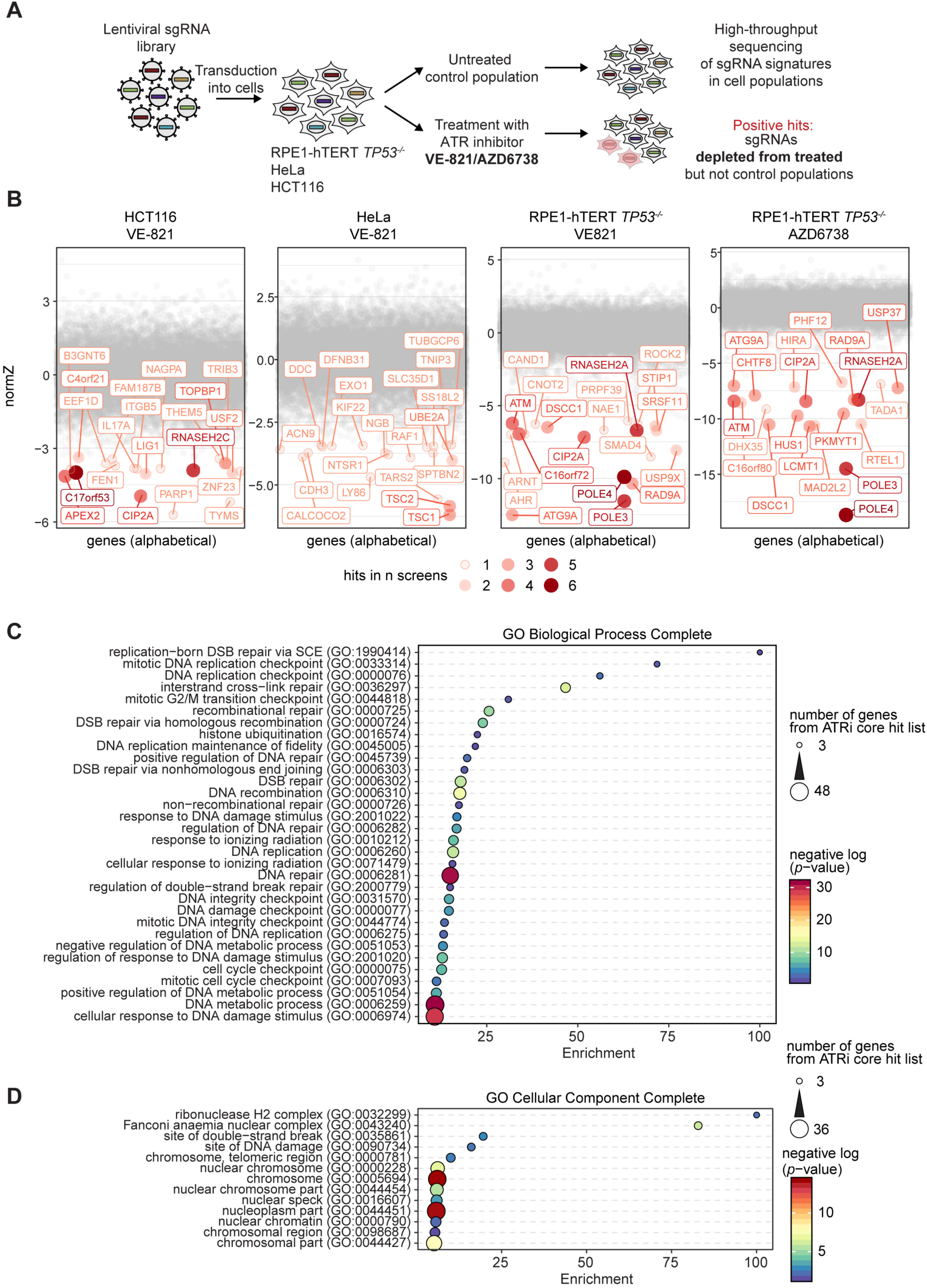
Identification of mutations that sensitize cells to ATR inhibition. a, Schematic of genome-wide CRISPR/Cas9 screen work flow. b, NormZ values were plotted against gene names in alphabetical order. For each screen, the genes with the 20 lowest NormZ values are labeled and coloured. 20 Genes with lowest NormZ values are labelled. Color, size and transparency of circles indicate number of screens (our datasets and datasets from Wang et al. 2018) in which the genes were hits (i.e. showed NormZ values < −2.5). c, Gene Ontology (GO) term enrichment analysis of Biological Process Complete terms (http://geneontology.org/page/go-enrichment-analysis) of 117 genes that were hits in at least 2 out of 7 screens using default settings. Shown are GO terms that are enriched at least 10-fold. Circle size indicates number of genes from 117-gene core set included in each GO term, color indicates negative log p-value and x-axis position indicates the fold enrichment compared to the whole genome reference set. d, GO term enrichment analysis of Cellular Component Complete terms as in c Shown are GO terms that are enriched at least 5-fold. DSB, DNA double strand break; SCE, sister chromatid exchange.

We initially screened Flag-Cas9 expressing clones of three cell types: HCT116 cells, derived from a colon carcinoma; HeLa cells, derived from a cervical carcinoma and a p53-mutated clone of RPE1 hTERT, which are telomerase-immortalized retinal pigment epithelial cells. These three cell lines were screened with the TKOv1 sgRNA library (Zimmermann et al. 2018) using VE-821 as the ATR inhibitor (Reaper et al. 2011). In the fourth screen, the library used was the newer generation TKOv3 (Hart et al. 2017) and AZD6738 was employed as the ATR inhibitor (Vendetti et al. 2015). The gene-level results can be found in Supplementary Table 1.

Using a hit-selection threshold based on *p*-values < 0.001, we found 32, 34 and 130 genes that promoted ATRi resistance in the HCT116, HeLa and RPE1-hTERT *TP53*^*-/-*^ cell lines, respectively using the TKOv1/VE-821 combination (Supplementary Table 1). In the RPE1-hTERT *TP53*^*-/-*^ cell line screened with TKOv3 and AZD6728, 88 hits were found and there was a good agreement with the two RPE1 screens (Fig 1B), with 41 common hits at *p*<0.001 (Supplementary Table 1). This good overlap suggests that both VE-821 and AZD6738 produce comparable phenotypes. In addition to these four screens, a recent publication also reported three CRISPR screens with AZD6738 as an ATR inhibitor in the MCF10A, HEK293 and HCT116 cell lines using the TKOv3 library (Wang et al. 2018). We re-analyzed this second set of screens using the newest version of drugZ (Wang et al. 2017) in order to provide a comparable set of data. We then combined the results of all seven screens and selected genes that were hits at a normalized z-score value (NormZ) less than −2.5 in at least two screens, which defined a set of high-confidence genes whose mutations cause ATR inhibitor sensitivity; this approach resulted in a “core” set of 117 genes. Gene ontology (GO) enrichment analyses (Fig 1CD) indicated that this set is highly enriched in GO terms associated with DNA replication, DNA repair and DNA damage checkpoint such as Replication-born DSB repair via SCE (GO:1990414), DNA replication checkpoint (GO:0000076) and Recombinational repair (GO:0000725) among the top enriched terms. Similarly, GO term analysis for Cellular Component yielded, Ribonuclease H2 complex (GO:0032299), the Fanconi Anemia nuclear complex (GO:0043240) and site of double-strand break (GO:0035861) as the top 3 enriched terms. Analysis of the gene set using the Reactome pathway database similarly identified highly connected pathways that revolve around DNA repair, DNA replication and cell cycle (Supplementary Fig 1).

In this core set, 11 genes were found as hits in at least 4 out of 7 screens indicating that they are likely to modulate the response to ATR inhibition independently of cellular context. These genes were *APEX2, ATM, ATRIP, C16orf72, C17orf53, KIAA1524* (also known as *CIP2A*), *POLE3, POLE4, RNASEH2A, RNASEH2B* and *RNASEH2C*. The sensitization of ATM-deficient cells to ATR inhibition had been described before (Reaper et al. 2011). Similarly, ATRIP is an activator of ATR (Zou and Elledge 2003) and we surmise that reducing ATR activity following *ATRIP* loss-of-function mutation sensitizes cells to ATR inhibition. In support of this possibility, ATR itself was a hit in RPE1-hTERT *TP53*^*-/-*^ cells/AZD6738 (*p* = 1.60 x 10^−7^), in the HEK293/AZD6738 (*p*= 0.0146) and HeLa/VE-821 cells (*p*= 0.0144; Supplementary Table 1). Drug sensitization by mutation of the drug target is a well-known phenomenon that has been harnessed to uncover drug targets in budding yeast (Giaever et al. 1999). The trimeric ribonuclease RNase H2 enzyme was recently described to promote resistance to ATR inhibitors (Wang et al. 2018). RNase H2 also promotes resistance to PARP inhibition, and RNase H2-deficient cells experience replication-associated DNA damage that depends on topoisomerase I (Zimmermann et al. 2018). The replication-associated DNA lesions caused by defective ribonucleotide excision repair in RNase H2-deficient cells may cause this observed vulnerability to ATRi. *APEX2* codes for the APE2 nuclease, which has been implicated in the regulation of ATR activity in *Xenopus laevis* cell-free extracts (Lin et al. 2018) and was recently found to be synthetic lethal with BRCA1 and BRCA2 deficiency (Mengwasser et al. 2019). These findings support the notion that the 117-gene core set identifies genetic determinants of the response to ATR inhibition.

To functionally validate the results, we selected 18 genes that were hits in the screens carried out in our laboratory (Supplementary Table 2). Of these, 15 out of 18 were part of the 117-ATRi core gene set. We undertook two-color competitive growth assays in which Cas9-expressing cells were transduced with lentiviral vectors that simultaneously express an sgRNA that targets a gene of interest (*GOI*) as well as GFP, or a control virus that expresses an sgRNA targeting *LacZ* and mCherry (Fig 2A). We carried out these assays first in RPE1-hTERT *TP53*^-/-^ cells. Two independent sgRNAs were tested per gene, and in some cases we monitored indel formation by TIDE analysis (Brinkman et al. 2014) to ensure formation of loss-of-function mutations (Supplementary Fig 2). The infected cell mixtures were grown in the presence or absence of VE-821 at doses of 2 and/or 4 μM, over the course of 15 d, and the proportions of GFP-and mCherry-expressing cells was determined at regular intervals by high-content microscopy. Out of this first set of analyses, the sgRNAs for 14 out of 18 genes clearly caused ATRi sensitivity (Fig 2B) whereas the sgRNAs targeting the remaining 4 genes (*NAE1, DPH1, DTYMK* and *PPP1R8*) produced inconclusive results because the sgRNAs themselves were highly cytotoxic in the absence of ATRi treatment (Supplementary Fig 3). However, as the set of 18 tested genes was not chosen at random, it is likely that the false positive rate might be slightly higher than 22% (4/18). Remarkably, nearly all the genes surveyed that were part of the core set (13/15) were successfully validated. Importantly, we validated the five (*APEX2, C17orf53, CIP2A, POLE3* and *POLE4*) of the 11 previously mentioned genes that were found in at least 4 out of 7 screens, highlighting their importance in the response to loss of ATR function.

**Figure 2.**
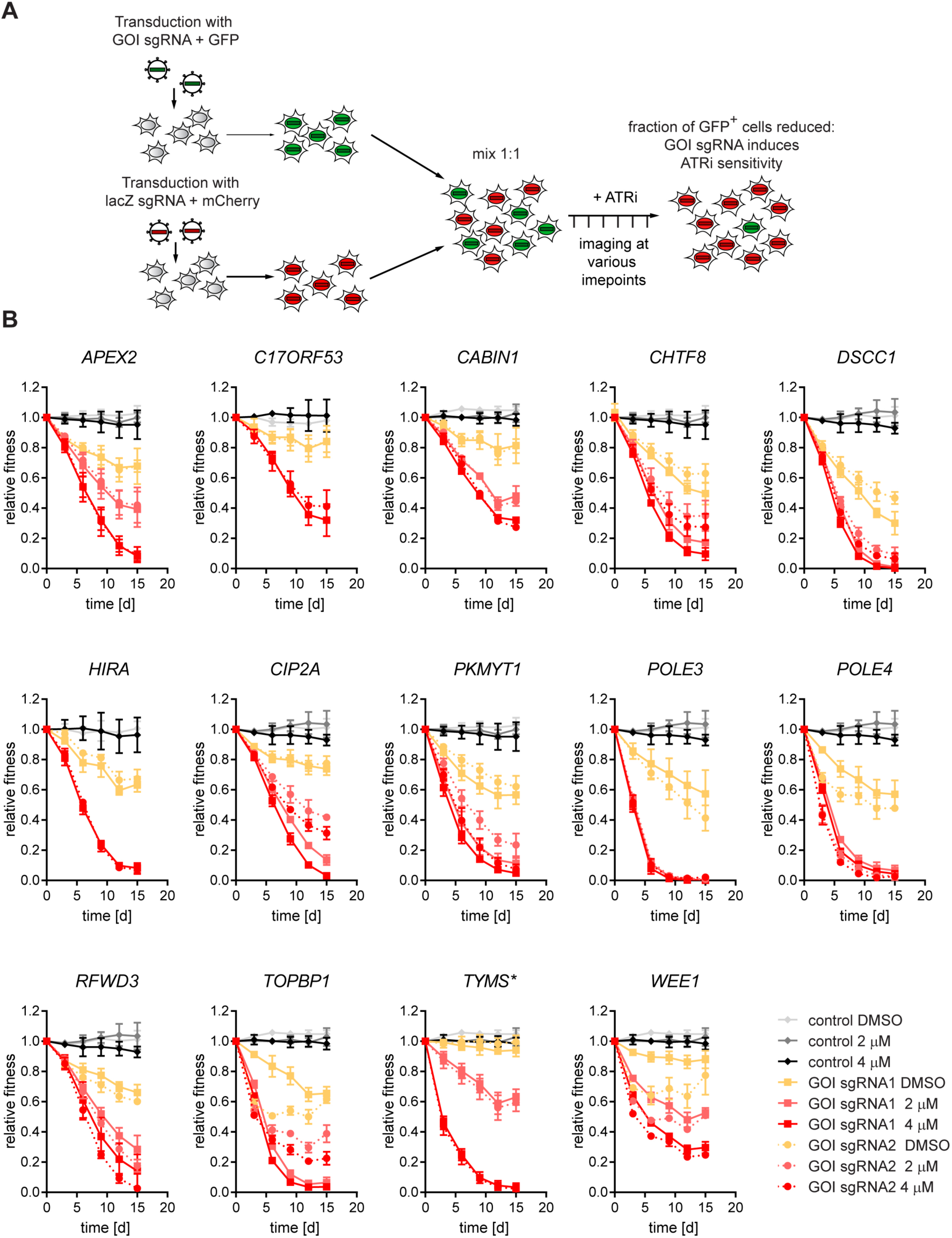
Hit validation using two sgRNAs for each gene of interest. a, Schematic showing work flow of two-color competitive growth assay. Cells were transduced with either an empty GFP vector (control) or vector with sgRNA targeting a gene of interest (GOI) coupled with GFP and an sgRNA targeting lacZ coupled with mCherry. GFP-and mCherry-expressing cells were mixed, treated or not with ATR inhibitor and population composition was followed over time. b, Two-color competitive growth assay results in RPE1-hTERT Flag-Cas9 *TP53*^*-/-*^ cells using either empty GFP vector control or one of two sgRNAs targeting a GOI coupled with GFP as well as a sgRNA targeting lacZ coupled with mCherry. Populations were treated with indicated concentrations of ATR inhibitor VE-821 or vehicle (DMSO) throughout the duration of the experiment. *HIRA*-and *C17orf53*-targeted cells were treated with only 0 or 4 µM VE-821. Plotted are the fraction of GFP-positive cells normalized to T0. Asterisks indicate genes that are not part of ATRi core gene set. Error bars represent standard deviation of three biologically independent experiments.

As a second stage of validation, we selected 8 genes (*APEX2, C17orf53, CABIN1, CIP2A, DSCC1, POLE4, TOPBP1, TYMS*) and assessed the ability of sgRNAs targeting them to engender sensitivity both to a second ATRi (AZD6738) and a second cell line (HCT116 cells). We found that 6 out of 8 genes promoted resistance to VE-821 and AZD6738 in both RPE1-hTERT *TP53*^*-/-*^ and HCT116 cells (Fig 3). The sgRNAs targeting *DSCC1* and *CABIN1* did not validate in HCT116 cells but we did not investigate further whether this was due to incomplete editing or whether it reflected biological differences between those cell lines.

**Figure 3.**
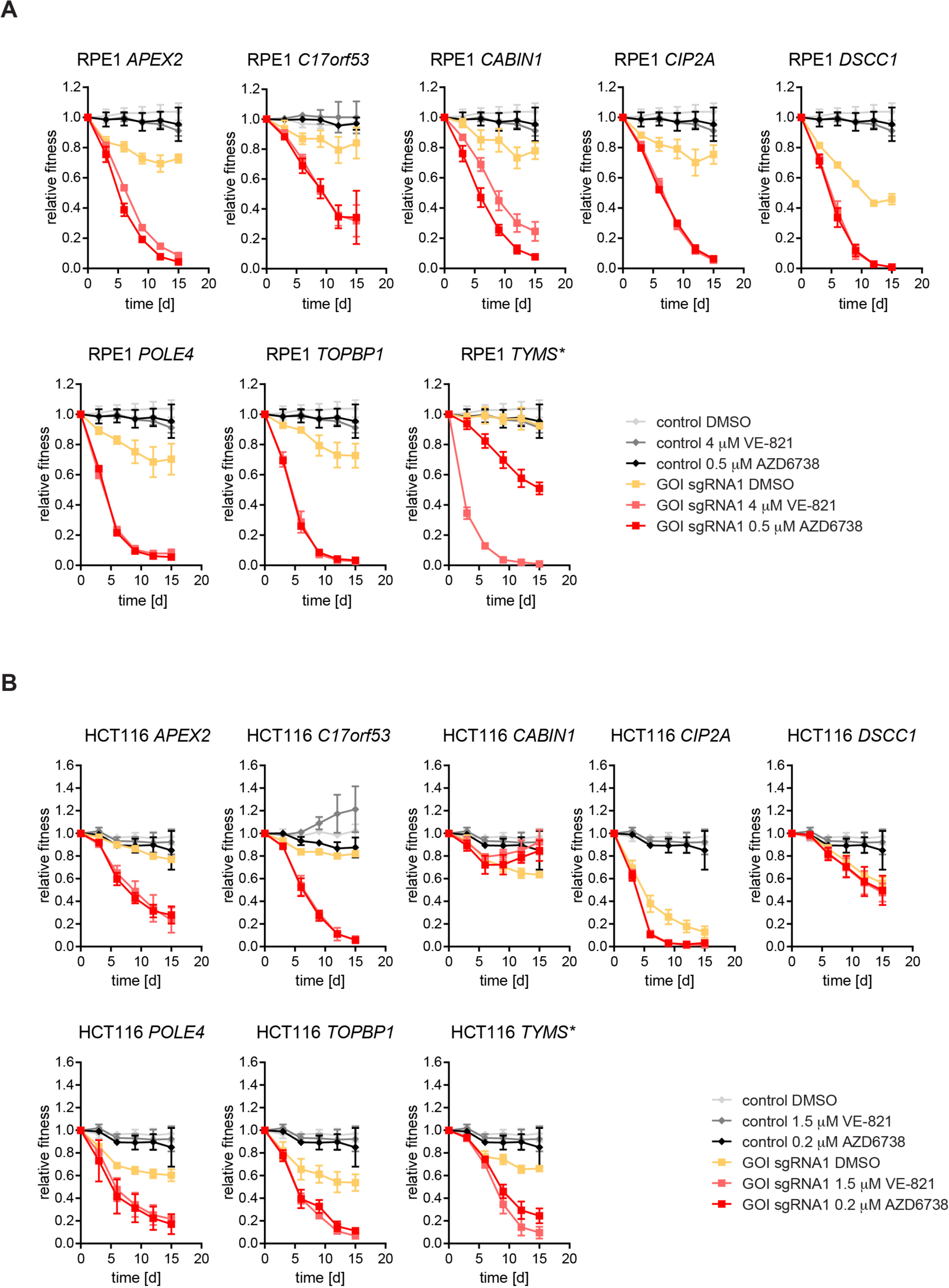
Hit validation using two ATR inhibitors and two cell lines. a, Results from two-color competitive growth assays using RPE1-hTERT Flag-Cas9 *TP53*^-/-^ cells and the indicated concentrations of ATRi (VE-821 or AZD6738) or vehicle (DMSO). b, Results from two-color competitive growth assays as in panel a, but using HCT116 Cas9 cells. Asterisks indicate genes that are not part of ATRi core gene set. Error bars represent standard deviation of three biologically independent experiments.

As a final stage of validation, we generated clonal loss-of-function mutations in the *APEX2, CIP2A, POLE3* and *POLE4* genes (Fig 4 and Supplementary Figs 4,5). We also added clones of *C16orf72* loss-of-function mutants as they were available in the laboratory (Fig 4 and Supplementary Fig 4). We assessed sensitivity to AZD6738 in clonogenic survival assays and observed that disruption of each of these genes caused hypersensitivity to ATR inhibition, with the mutations in the *POLE3* and *POLE4* genes causing the greatest sensitization to ATR inhibition (Fig 4C,D), in line with the results obtained in the competitive growth assay (Figs 2,3).

**Figure 4.**
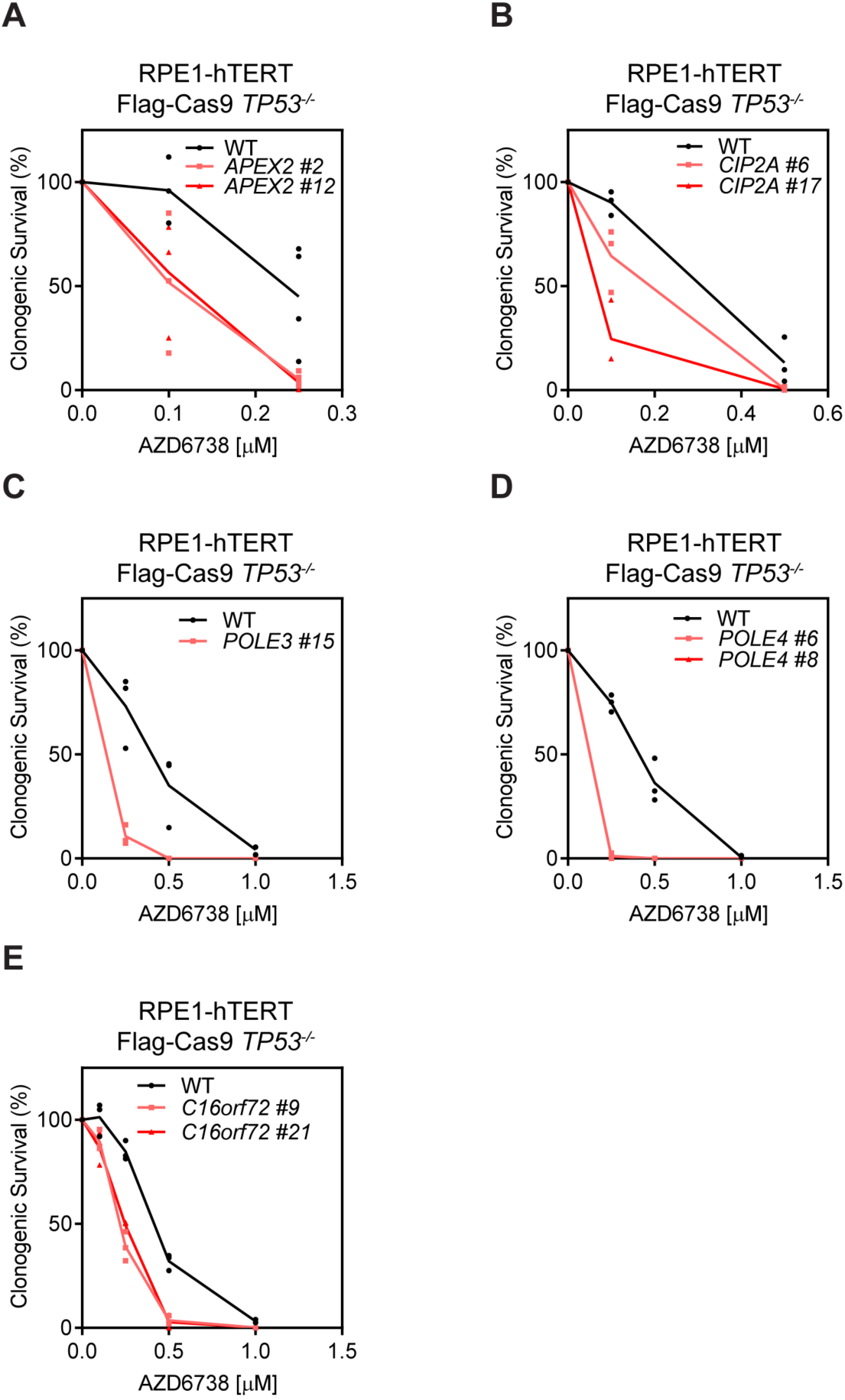
Clonal KO cell lines of *APEX2, CIP2A, POLE3, POLE4* and *C16ORF72* are sensitive to ATRi. a, Clonogenic survival of RPE1-hTERT Flag-Cas9 *TP53*^-/-^ (WT) and two RPE1-hTERT Flag-Cas9 *TP53*^-/-^ *APEX2*^-/-^ clones treated with indicated concentrations of ATR inhibitor AZD6738. b, as in a using two *CIP2A*^*-/-*^ clones. c, as in a using a *POLE3*^*-/-*^ clone. d, as in a using two *POLE4*^*-/-*^ clones. e, as in a using two *C16orf72*^-/-^ clones. Data are from three biologically independent experiments.

The remarkable hypersensitivity of POLE3/POLE4-deficient cells to ATR inhibitors was intriguing in light of the recent characterization of this protein complex in chromatin maintenance and within the DNA Pol ε holoenzyme (Bellelli et al. 2018a; Bellelli et al. 2018b; Goswami et al. 2018; Yu et al. 2018). The POLE3/4 subunits form a histone-fold complex that is flexibly tethered to the core subunits of Pol ε but is not essential for DNA polymerization (Goswami et al. 2018). POLE3/4 acts as a histone H3-H4 chaperone that ensures symmetric histone deposition during DNA replication (Bellelli et al. 2018a; Yu et al. 2018). *Pole4*^*-/-*^ mice are viable and show evidence of replication stress despite the fact that *POLE4*^*-/-*^ cells have normal activation of the ATR pathway in response to a campthotecin or hydroxyurea challenge (Fig 5A) (Bellelli et al. 2018b). We tested whether the ATRi sensitivity of *POLE3*^*-/-*^ cells was due to defective histone deposition by complementing *POLE3*^-/-^ knockout cells with a vector expressing a variant POLE3 with a C-terminal deletion, POLE3ΔC, which disrupts the histone deposition function of the POLE3/POLE4 dimer (Bellelli et al. 2018a). Expression of wild type POLE3 or POLE3ΔC fully restored ATRi resistance suggesting that histone deposition by this complex is unlikely to be involved in the normal cellular resistance to ATR inhibitors (Fig 5B,C and Supplementary Figure 6).

**Figure 5.**
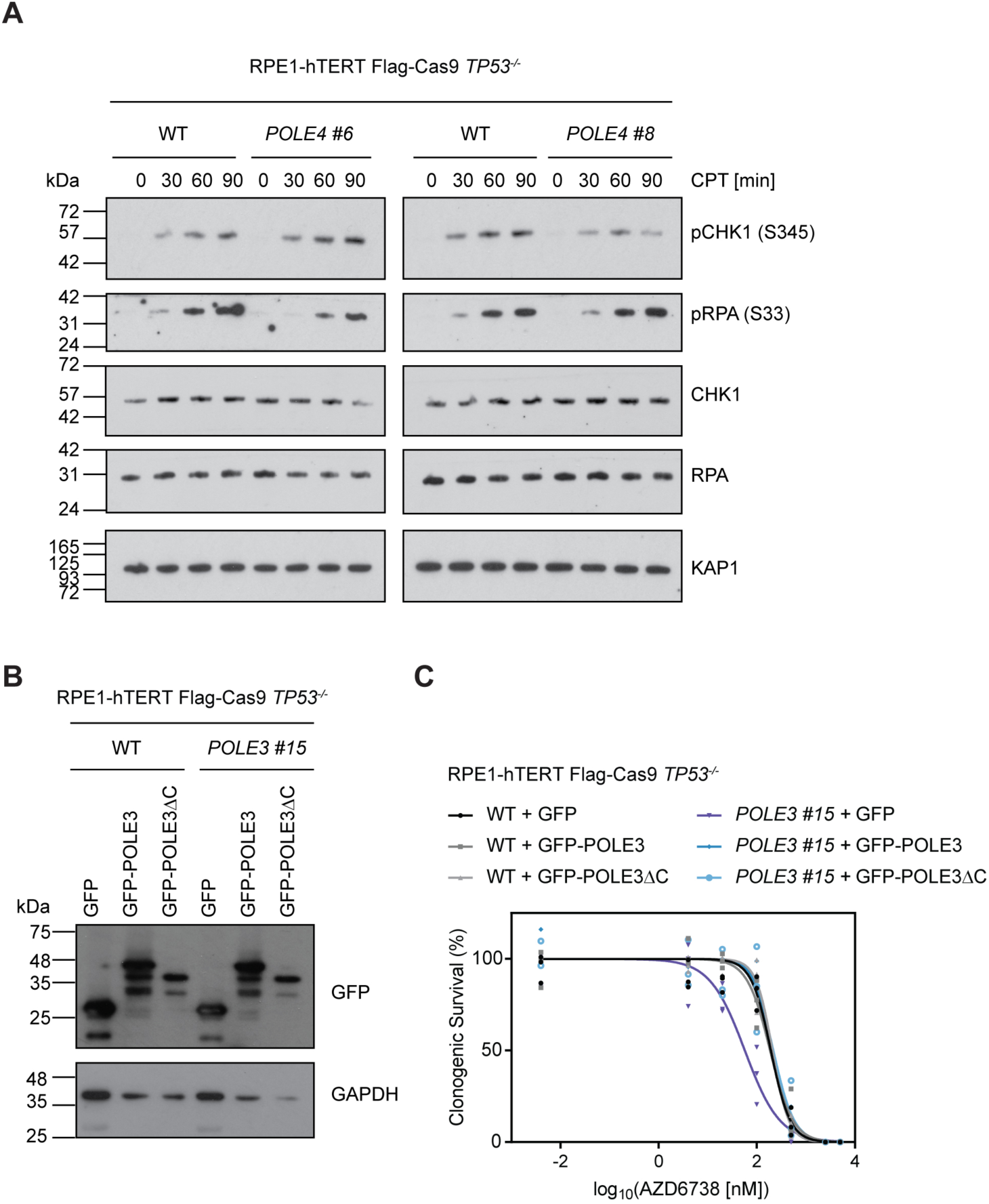
ATRi sensitivity in *POLE3*^-/-^ or *POLE4*^*-/-*^ cells is not caused by defective ATR signalling or histone deposition. a, Whole cell extracts from wild type RPE1-hTERT Flag-Cas9 *TP53*^*-/-*^ (WT) or the indicated *POLE4*^*-/-*^ clones treated with 1 μM camptothecin (CPT) were used for immunoblotting with indicated antibodies. pCHK1 and pRPA refer to phosphorylated proteins; brackets indicate modified amino acid residues. KAP1 served as loading control. b, Whole cell extracts from WT or the indicated *POLE3*^*-/-*^ clone expressing GFP, GFP-POLE3 or GFP-POLE3ΔC were used for immunoblotting with indicated antibodies. GAPDH served as loading control. c, Clonogenic survival of WT or the indicated *POLE3*^*-/-*^ clone expressing GFP, GFP-POLE3 or GFP-POLE3ΔC treated with indicated concentrations of ATR inhibitor AZD6738. Data are from three biologically independent experiments.

In summary, our mapping of gene mutations that cause sensitivity to ATRi provides an unbiased view of the genetic architecture of the ATR-dependent control of genome integrity. We contend that this dataset will be a valuable tool for both clinical and biological researchers. Beyond revealing potential new biomarkers of the ATRi response, this list is rich in potential new avenues for study. For example, there are many genes in this list that have never been characterized in-depth for a role in genome integrity. A prime candidate is *C4orf21*, also known as *ZGRF1*, which encodes a protein containing a GRF-type zinc finger, a domain found in bona fide DNA repair proteins such as TOPIIIα, APE2 and the NEIL3 glycosylase. In some other cases, the inclusion of a gene in this list cements the potential role of their encoded protein as a modulator of genome integrity. An example is DHX9, which is an RNA helicase that promotes R-loop formation and impedes DNA replication (Chakraborty et al. 2018). We also note that this core gene set is unlikely to be exhaustive because fitness screens necessitate that the sgRNAs have some level of representation in the control cell population and therefore can miss cell-essential genes. Given that DNA replication is an essential process associated with the ATR pathway, it is likely that a comprehensive list of vulnerabilities to ATR inhibition will necessitate other approaches such as phenotypic screens or CRISPR interference. Finally, our finding that mutations in *POLE3*/*POLE4* cause hyper-sensitivity to ATRi is particularly intriguing and forms the basis of ongoing studies. Our data indicates that the sensitivity imparted by the loss of POLE3/POLE4 is not due to defective histone deposition, suggesting it is rather caused by a defect in DNA synthesis that remains to be uncovered.

## Materials and Methods

### Plasmids

DNA sequences corresponding to sgRNAs were cloned into LentiGuide-Puro (Addgene: 52963) or a modified form of LentiCRISPRv2 (Addgene: 52961) in which the sequence encoding Cas9 was replaced by that for NLS-tagged GFP or mCherry using AgeI and BamHI (designated as LentiGuide-NLS-GFP or –mCherry), as described (Sanjana et al. 2014; Shalem et al. 2014). GFP-POLE3 full length and GFP-POLE3 with amino acid residues 113-140 deleted (GFP-POLE3ΔC) were cloned between NheI and AgeI restriction sites of pCW57.1 (Addgene: 41393),

### Cell lines and gene editing

293T cells were obtained from ATCC. HeLa Flag–Cas9 and RPE1-hTERT Flag–Cas9 *TP53*^*-/-*^ were published earlier (Zimmermann et al. 2018) and HCT116 Flag-Cas9 cells (Hart et al. 2015) were a kind gift from Jason Moffat. RPE1-hTERT Flag-Cas9 *TP53*^*-/-*^ were grown in Dulbecco’s Modified Eagle Medium (DMEM; Gibco /Wisent) supplemented with 10% fetal bovine serum (FBS; Wisent), 200 mM GlutaMAX, 1× non-essential amino acids (both Gibco) and 100 U ml^-1^ penicillin and 100 µg ml^-1^ streptomycin (Pen/Strep; Wisent/Gibco). HeLa Flag-Cas9 and 293T cells were cultured in DMEM supplemented with 10% FBS and Pen/Strep. HCT116 Flag-Cas9 cells were cultured in McCoy’s 5A medium (Gibco) supplemented with 10% FBS and Pen/Strep. Cell lines stably expressing Flag-Cas9 were maintained in the presence of 2 µg/mL blasticidin.

Lentiviral particles were produced in 293T cells by co-transfection of the targeting vector with plasmids expressing VSV-G, RRE and REV using TransIT LT-1 transfection reagent (Mirus). Viral transductions were performed in the presence of 4-8 µg/mL polybrene (Sigma-Aldrich) at an MOI < 1. Transduced RPE1-hTERT Flag-Cas9 *TP53*^*-/-*^ and HCT116 Cas9 cells were selected by culturing in the presence of 15-20 or 2 µg/mL puromycin, respectively.

*APEX2, CIP2A, POLE3 and POLE4* gene knockouts were generated in RPE1-hTERT Flag-Cas9 *TP53*^*-/-*^ cells by electroporation of LentiGuide-Puro or LentiGuide-NLS-GFP vectors using an Amaxa II Nucleofector (Lonza). *C16orf72* gene knockout clones were generated in RPE1-hTERT Flag-Cas9 *TP53*^*-/-*^ cells by transfecting sgRNA/Cas9 ribonucleoprotein complex using Lipofectamine CRISPRMAX Cas9 Transfection Reagent (Invitrogen). After 24 h transfection, cells were expanded, followed by single clone isolation. For sgRNA sequences employed see Supplementary Table 3; *APEX2 #12*: *APEX2*sgRNA1, *APEX2 #2*: *APEX2*sgRNA2, *C16orf72 #9* and *C16orf72 #21*: C16orf72sgRNA1, *CIP2A #6* and *CIP2A #17*: *CIP2A*sg RNA2, *POLE3 #15*: *POLE3*sgRNA1, *POLE4 #6* and *POLE4 #8*: *POLE4*sgRNA1. 24 h following transfection, cells were selected for 24-48 h with 15-20 µg/mL puromycin, followed by single clone isolation. Gene mutations were further confirmed by PCR amplification, DNA sequencing and TIDE analysis (Brinkman et al. 2014). For primers used for genomic PCR, see Supplementary Table 4. Loss of gene expression was further confirmed either by immunoblotting to assess protein levels if antibodies were available (see below) or by RT-qPCR to assess mRNA levels using GAPDH for normalization. Taqman assays employed were GAPDH (Hs99999905_m1) and APEX2 (Hs00205565_m1) from Thermo Fisher Scientific.

RPE1-hTERT Flag-Cas9 *TP53*^*-/-*^ (WT) or *POLE3*^*-/-*^ #15 cells expressing GFP, GFP-POLE3 or GFP-POLE3ΔC were generated by transduction with lentiviral particles of pCW57.1-derived GFP, -GFP-POLE3 or - GFP-POLE3ΔC constructs and subsequent selection with 20ug/mL Puromycin for 48h. Cells were maintained in the presence of 5 ug/mL puromycin and 1ug/mL doxycycline.

### Antibodies, siRNAs and drugs

The following antibodies were used in this study at the indicated dilutions: anti-CIP2A (CST #14805; 1:1000), anti-GAPDH (Sigma-Aldrich G9545; 1:20000), anti-GFP (gift from Laurence Pelletier, 1:10000), anti-KAP1 (Bethyl A300-274A, 1:5000), anti-POLE3 (Bethyl A301-245A-1; 1:2000), anti-POLE4 (Abcam ab220695; 1:200), anti-Tubulin (Millipore CP06, 1:2000), anti-pCHK1 (S345) (Cell Signaling #2348, 1:1000), anti-CHK1 (Santa Cruz sc8408, 1:500), anti-pRPA32 (S33) (Bethyl A300-246A-3, 1:20,000), anti-RPA32 (Abcam ab2175, 1:500). The following secondary antibodies for immunoblotting were used in this study: peroxidase-conjugated AffiniPure Bovine Anti-Goat IgG (Jackson Immuno Research 805-035-180) and peroxidase-conjugated Sheep Anti Mouse IgG (GE Healthcare NA931V). All peroxidase-conjugated secondary antibodies were used at a dilution of 1:5000. Protein bands were detected using the SuperSignal West Pico enhanced chemiluminescence reagent (Thermo Fisher Scientific). The following siRNAs from Dharamacon were used in this study: Control, siGENOME Non-targeting Pool #2 (D-001206-14-05); POLE3, siGENOME SMARTpool (M-008460-01-0005); POLE4, siGENOME SMARTpool (M-009850-01-0005); APEX2, siGENOME SMARTpool (M-013730-00-0005). ATR inhibitors VE-821 and AZD6738 were purchased from SelleckChem.

### CRISPR/Cas9 screens

RPE1-hTERT Flag–Cas9 *TP53*^*-/-*^, HeLa Flag-Cas9 and HCT116 Flag-Cas9-expressing cells were transduced with the lentiviral TKOv1 library at a low MOI (∼0.35) and puromycin-containing media was added the next day to select for transductants. Selection was continued until 72 h post-transduction, which was considered the initial time point, t0. To identify VE-821 sensitizers, the negative-selection screen was performed by subculturing at days 3 and 6 (t3 and t6), at which point the cultures were split into two populations. One was left untreated and to the other a dose of VE-821 amounting to 20% of the lethal dose (LD_20_) (HeLa Flag-Cas9, 1.5 µM; HCT116 Flag-Cas9 1.5 µM; RPE1-hTERT Flag-Cas9 *TP53*^*-/-*^, 4 µM) was added. Cells were grown with or without VE-821 until t18 and subcultured every three days. Sample cell pellets were frozen at t18 for genomic DNA (gDNA) isolation. Screens were performed in technical triplicates and a library coverage of ≥200 cells per sgRNA was maintained at every step. The AZD6738 screen was performed at a concentration of 0.5 µM AZD6738 using the TKOv3 library (Hart et al. 2017) in technical duplicates and a library coverage of ≥ 375 cells per sgRNA was maintained at every step. Genomic DNA from cell pellets was isolated using the QIAamp Blood Maxi Kit (Qiagen) and genome-integrated sgRNA sequences were amplified by PCR using the KAPA HiFi HotStart ReadyMix (Kapa Biosystems). i5 and i7 multiplexing barcodes were added in a second round of PCR and final gel-purified products were sequenced on Illumina NextSeq500 or HiSeq2500 systems to determine sgRNA representation in each sample. sgRNA sequence read counts (Supplementary Table 5) were obtained using MaGeck (Li et al. 2014). drugZ (Wang et al. 2017) was used to identify gene knockouts which were depleted from ATRi-treated t18 populations but not depleted from untreated cells.

### Two-color competitive growth assay

Cells were transduced with either virus particles of NLS-mCherry LacZ-sgRNA or NLS-GFP GOI-sgRNA. 24h after transduction transduced cells were selected using 15-20 µg/mL puromycin for 48 h. At this time mCherry-and GFP-expressing cells were mixed 1:1 (2,500 cells + 2,500 cells) and plated in a 12-well format. Cells were imaged for GFP-and mCherry signals 24h after initial plating (t=0) and ATR inhibitors were added subsequently (RPE1-hTERT Flag-Cas9 *TP53*^*-/-*^: 2µM/4 µM VE-821 or 0.5 µM AZD6738; HCT116 Flag-Cas9: 1.5µM VE-821 and 0.2 µM AZD6738). During the course of the experiment, cells were subcultured when near-confluency was reached and imaged on days 3, 6, 9, 12, and 15. An InCell Analyzer system (GE Healthcare Life Sciences) equipped with a 4X objective was used for imaging. Segmentation and counting of the number of GFP-positive and mCherry-positive cells was performed using an Acapella script (PerkinElmer). Efficiency of indel formation for a subset of sgRNAs was analysed by performing PCR amplification and sequencing of the region surrounding the sgRNA target sequence and TIDE analysis on DNA isolated from GFP-expressing cells 9 days post-transduction.

### Clonogenic survival assays

RPE1-hTERT Flag-Cas9 *TP53*^*-/-*^ cells were seeded in 10-cm dishes (WT: 500 or 1000 cells; *APEX2*^-/-^: 1000-1500 cells; *CIP2A*^*-/-*^: 2000 cells; *POLE3*^*-/-*^: 1000 cells *POLE4*^-/-^: 1000 cells) in the presence AZD6738 or left untreated. For WT or POLE3*-/-*cells transduced with pCW57.1-GFP/GFP-POLE3/GFP-POLE3ΔC 1000 cells were seeded in 10 cm dishes in the presence of 1μg/mL doxycycline and in the presence or absence of AZD6738. Doxycycline-containing medium +/-AZD6738 was refreshed every three days. After 11-14 days, colonies were stained with crystal violet solution (0.4 % (w/v) crystal violet, 20% methanol) and counted manually. Relative survival was calculated for the drug treatments by setting the number of colonies in non-treated controls at 100%.

### Incucyte assay for ATRi sensitivity

RPE1-hTERT Flag-Cas9 *TP53*^*-/-*^ (WT) or *POLE3*^*-/-*^ cells were seeded into a 12-well plate (2500 cells per well) in the presence of 1μg/mL doxycycline. After 24 h medium containing 1μg/ml doxycycline and AZD6738 to the desired final concentration was added. Every six hours 25 images per well were acquired using an Incucyte S3 Live Cell Analysis System (Sartorius) and analyzed for percentage confluency using Incucyte S3 2018A software (Sartorius). Doxycycline-containing medium +/-AZD6738 was refreshed after three days. Percentage confluency at the 138 h time point was used to calculate relative survival by setting the percentage confluency in non-treated controls at 100%.

## Acknowledgments

We thank Rachel Szilard for the critical reading of this manuscript. Jason Moffat for TKO libraries and cell lines, Laurence Pelletier for sharing his GFP antibody and Kin Chan of the LTRI NBCC for sequencing. NH is supported by a Human Frontier Science Program long-term Fellowship. TH is supported by MD Anderson Cancer Center Support Grant P30 CA016672 and the Cancer Prevention Research Institute of Texas (CPRIT/RR160032). DD is a Canada Research Chair (Tier I). DD is funded by CIHR grant FDN143343 and Canadian Cancer Society (CCS grant #705644).

---

**Supplementary Figure 1.**
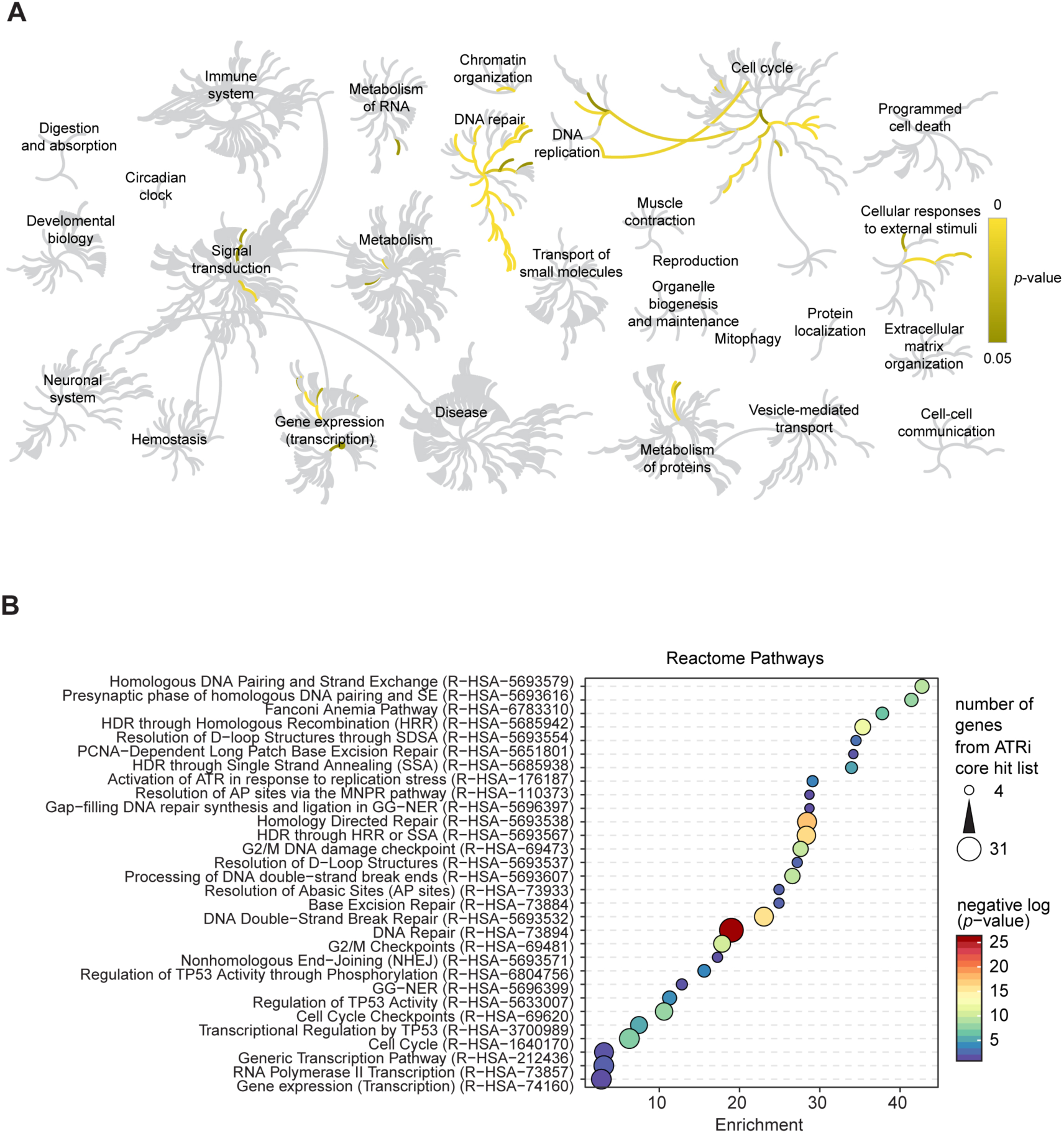
Reactome pathways enrichment analysis. a, The 117-gene ATRi core gene set was analyzed for enrichment of Reactome pathways (https://reactome.org). Shown is Reactome pathway browser with significantly enriched pathways highlighted in yellow. Color bar indicates *p*-value. b, Results of enrichment analysis of Reactome pathways (http://geneontology.org/page/go-enrichment-analysis) as in Figure 1c and d. Shown are all enriched Reactome pathways except unclassified genes. HDR, homology-directed repair; SDSA, synthesis-dependent strand annealing; MNPR, multiple-nucleotide patch replacement; GG-NER, global genome nucleotide excision repair; HRR, homologous recombination repair, SSA, single strand annealing.

**Supplementary Figure 2.**
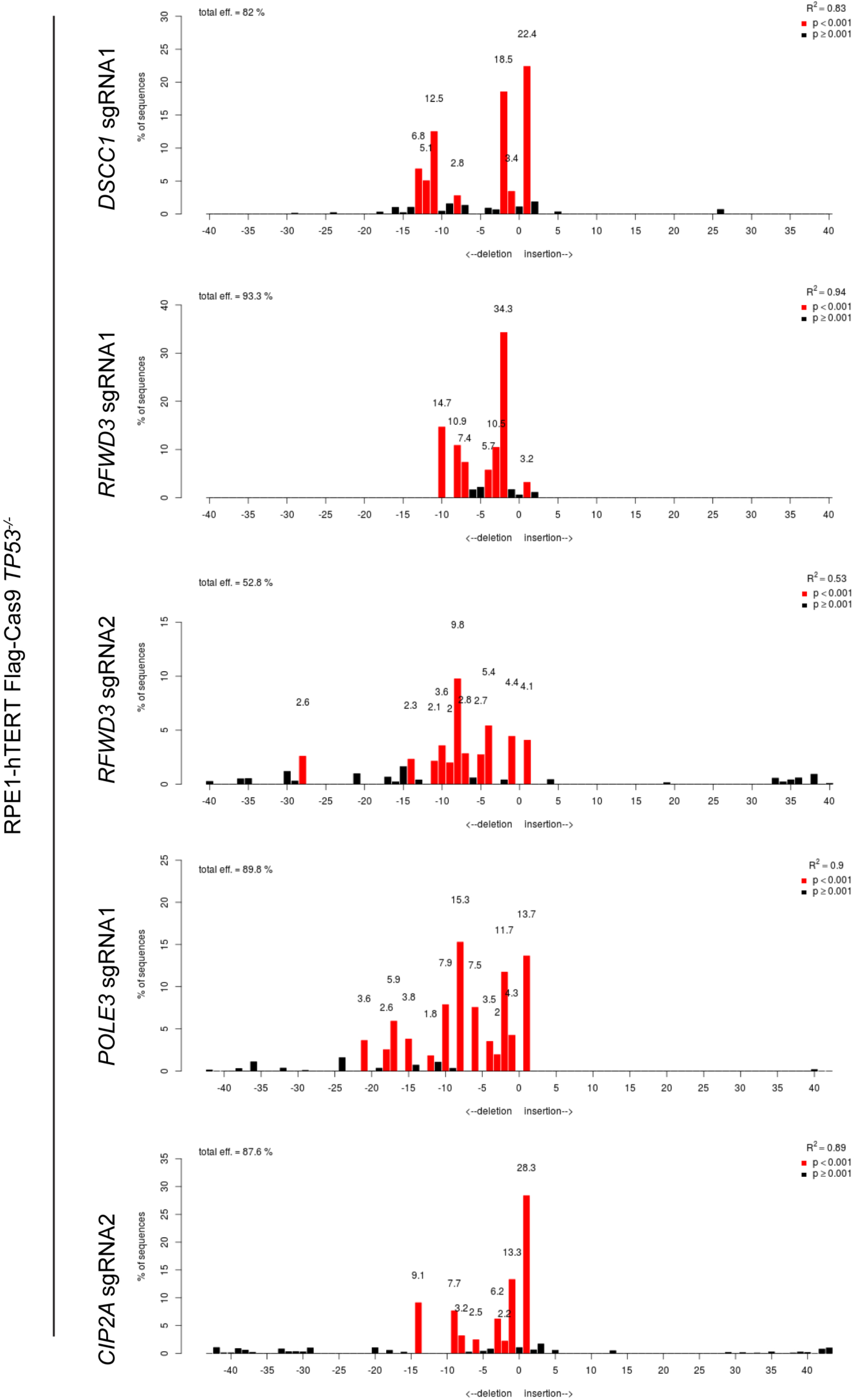
Editing efficiency of selected sgRNAs. Genomic DNA was isolated from RPE1-hTERT Flag-Cas9 *TP53*^*-/-*^ cells transduced with indicated sgRNAs (for sequences see Supplementary Table 3). The region around the sgRNA targeting site was amplified by PCR and sequenced. Results from TIDE analysis of obtained sequences are shown.

**Supplementary Figure 3.**
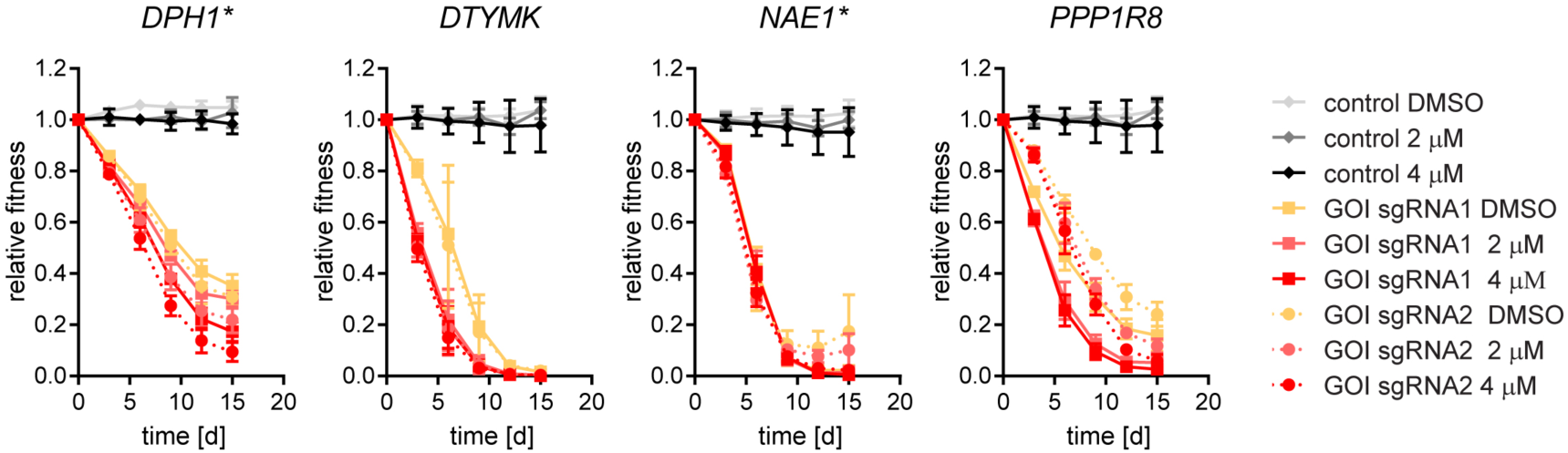
Hit validation using two sgRNAs. a, Results of two-color competitive growth assays. RPE1-hTERT Flag-Cas9 *TP53*^*-/-*^ cells were transduced with either an empty GFP vector (control) or one of two sgRNAs targeting *DPH1, DTYMK, NAE1* or *PPP1R8* coupled with GFP as well as an sgRNA targeting lacZ coupled with mCherry and treated with indicated concentrations of VE-821. Asterisks indicate genes that are not part of ATRi core gene set. Error bars represent standard deviation of three biologically independent experiments.

**Supplementary Figure 4.**
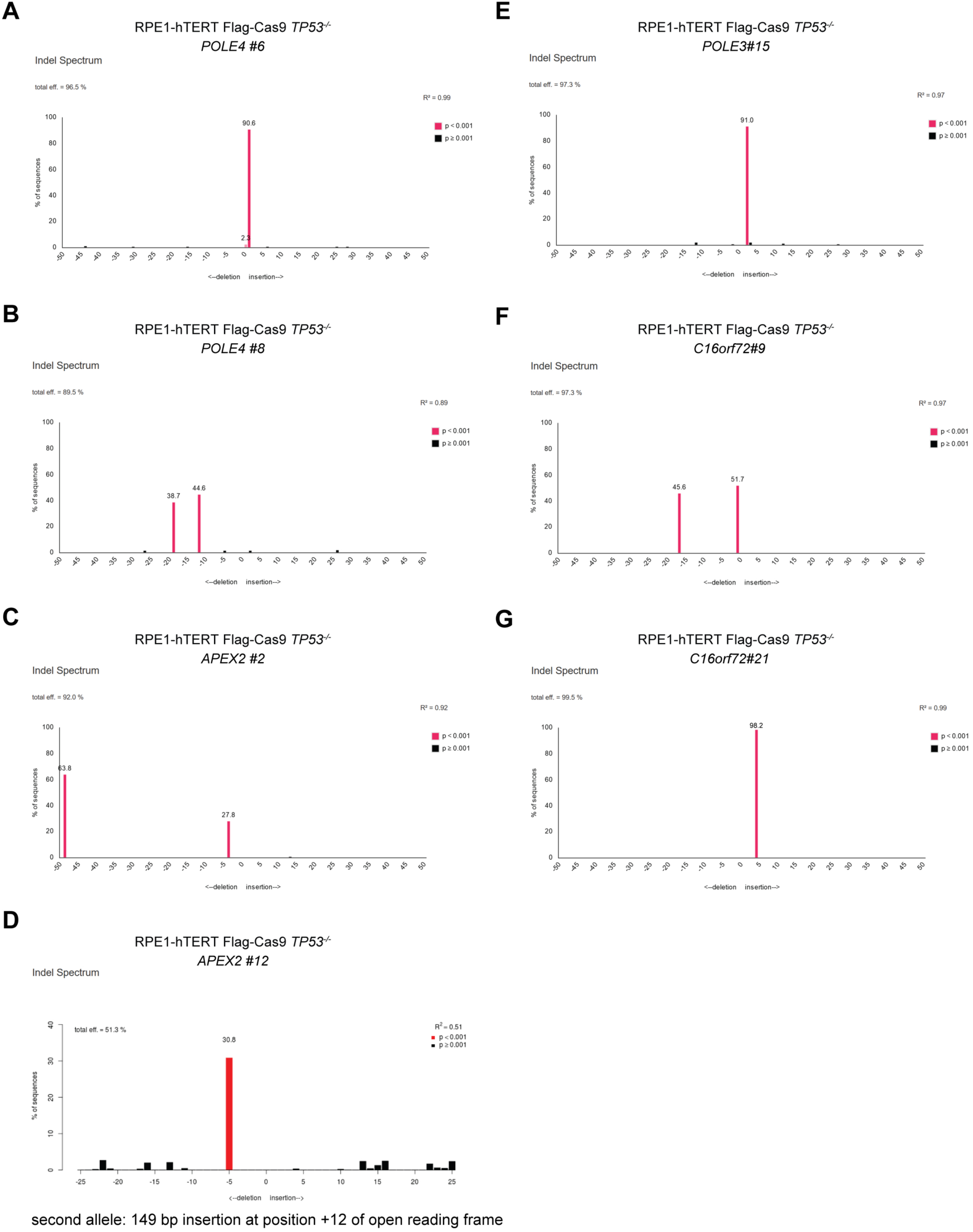
Validation of clonal KO cell lines by TIDE analysis. a-g, TIDE analysis of sequences from indicated clones surrounding the targeting sites of sgRNAs used to create gene knockouts.

**Supplementary Figure 5.**
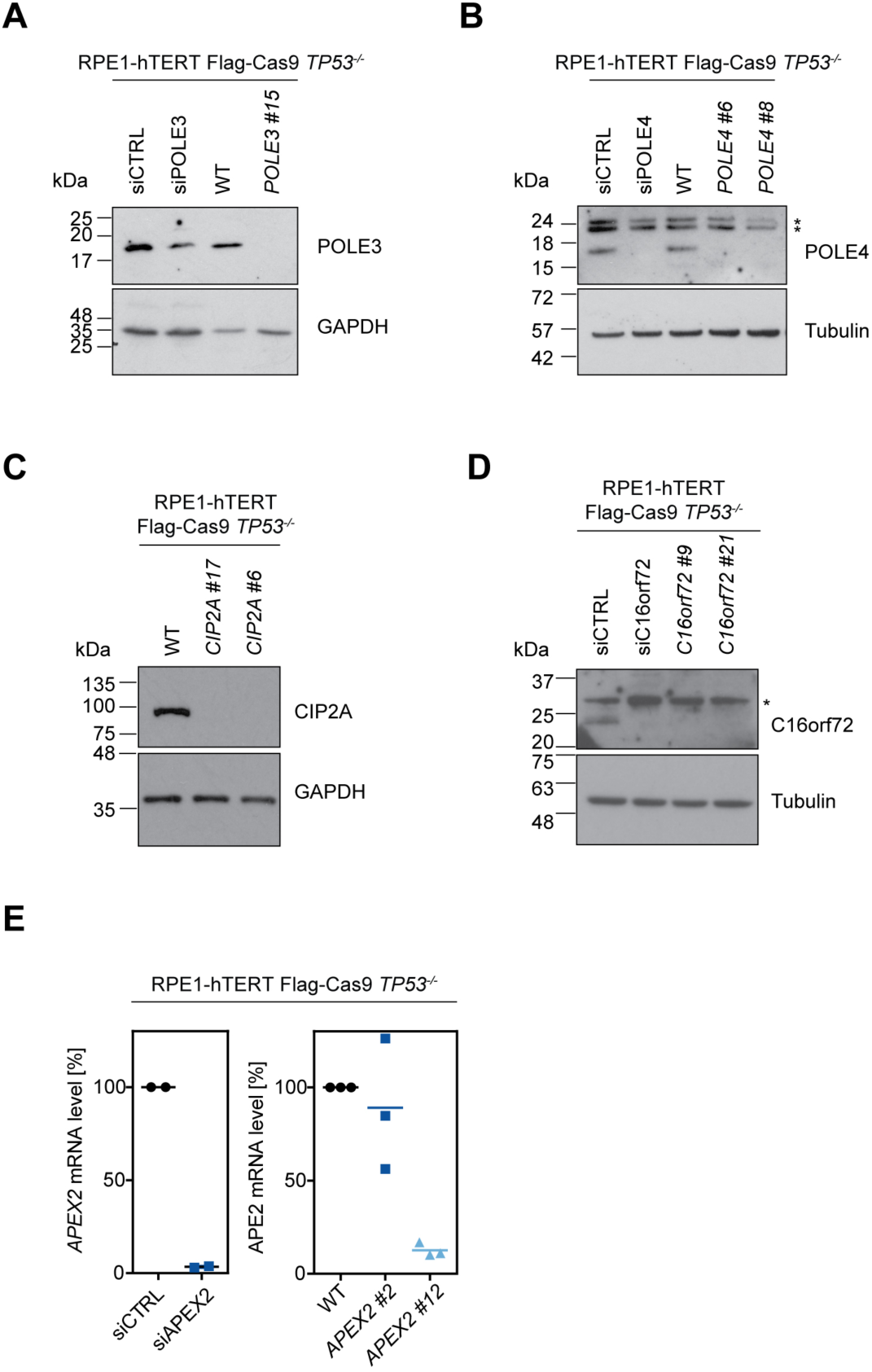
Validation of clonal KO cell lines by immunoblotting or qPCR. a-c, Immunoblotting to assess loss of protein expression in clonal knockout cell lines. siRNA-mediated knockdown was used to control for antibody specificity in the case of POLE3 and POLE4 antibodies. Antibodies targeting alpha-tubulin (Tubulin) or GAPDH were used as loading controls. Numbers indicate molecular mass in kDa. Asterisks indicate unspecific bands. d, mRNA level analysis of *APEX2* after siRNA mediated *APEX2* knockdown as assay control and in *APEX2*^-/-^ clones. Clone #2 showed mRNA levels similar to parental (WT) cells but has frameshifting mutations (see panel D) and was sensitive to ATR inhibitor treatment (see Figure 4).

**Supplementary Figure 6.**
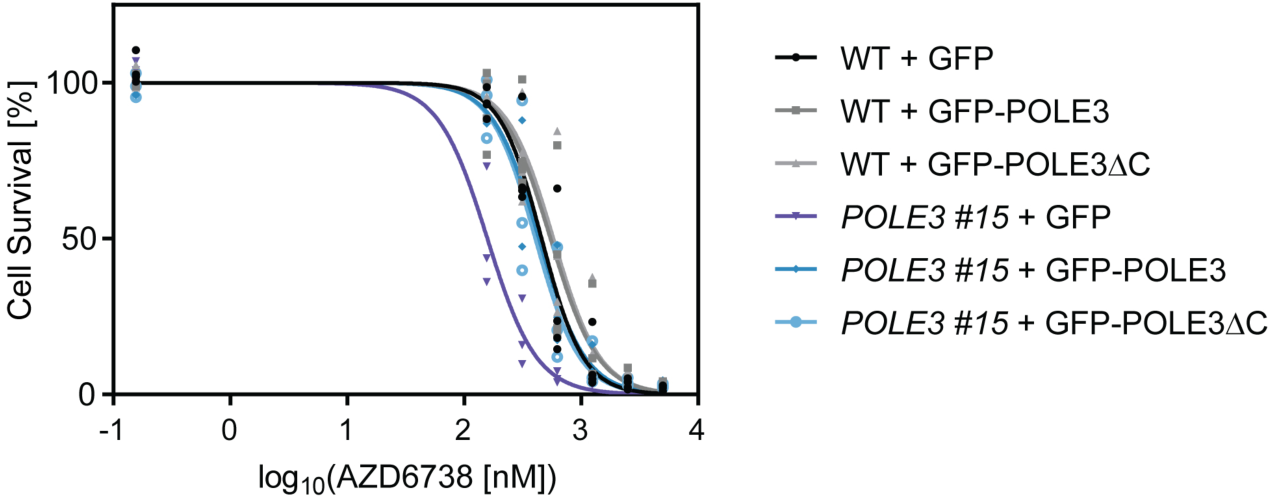
POLE3ΔC mutant is not sensitive to ATRi by in an Incucyte assay. Cell survival of RPE1-hTERT Flag-Cas9 *TP53*^*-/-*^ (WT) or the indicated *POLE3*^-/-^ clone expressing GFP, GFP-POLE3 or GFP-POLE3ΔC treated with indicated concentrations of ATR inhibitor (AZD6738) was determined by monitoring growth in an Incucyte instrument. Data are from three biologically independent experiments.

